# Novel approaches to label the surface of *S. aureus* with DBCO for click chemistry-mediated deposition of sensitive cargo

**DOI:** 10.1101/2024.12.18.629304

**Authors:** Tsvetelina H. Baryakova, Laura Segatori, Kevin J. McHugh

## Abstract

The strain-promoted alkyne-azide cycloaddition (SPAAC) reaction can be used to modify the surface of bacteria for a variety of applications, including drug delivery, biosensing, and imaging. This is usually accomplished by first installing a small azide group within the peptidoglycan and then delivering exogenous cargo (e.g., a protein or nanoparticle) modified with a cyclooctyne group, such as dibenzocyclooctyne (DBCO), for *in situ* conjugation. However, DBCO is comparatively bulky and hydrophobic, increasing the propensity for some payloads to aggregate. In this study, we sought to invert this paradigm by exploring two novel strategies for incorporating DBCO into the peptidoglycan of *Staphylococcus aureus* and compared them to an established approach using DBCO-vancomycin. We demonstrate that DBCO-modified small molecules belonging to all three classes – a sortase peptide substrate (LPETG), two D-alanine derivatives, and vancomycin – can selectively label the *S. aureus* surface to varying degrees. In contrast to DBCO-vancomycin, the DBCO-D-alanine variants do not adversely affect the growth of *S. aureus* or lead to off-target labeling or toxicity in HEK293T cells, even at high concentrations. Finally, we show that, unlike IgG3-Fc labeled with DBCO groups, IgG3-Fc labeled with azide groups is stable (i.e., remains water-soluble) under normal storage conditions, retains its ability to bind the immune receptor CD64, and can be successfully attached to the surface of DBCO-modified *S. aureus*. We believe the labeling strategies explored herein will expand the paradigm of specific, nontoxic SPAAC-mediated labeling of the surface of *S. aureus* and other gram-positive bacteria, opening the door for new applications using azido-modified cargo.

**GRAPHICAL ABSTRACT:** 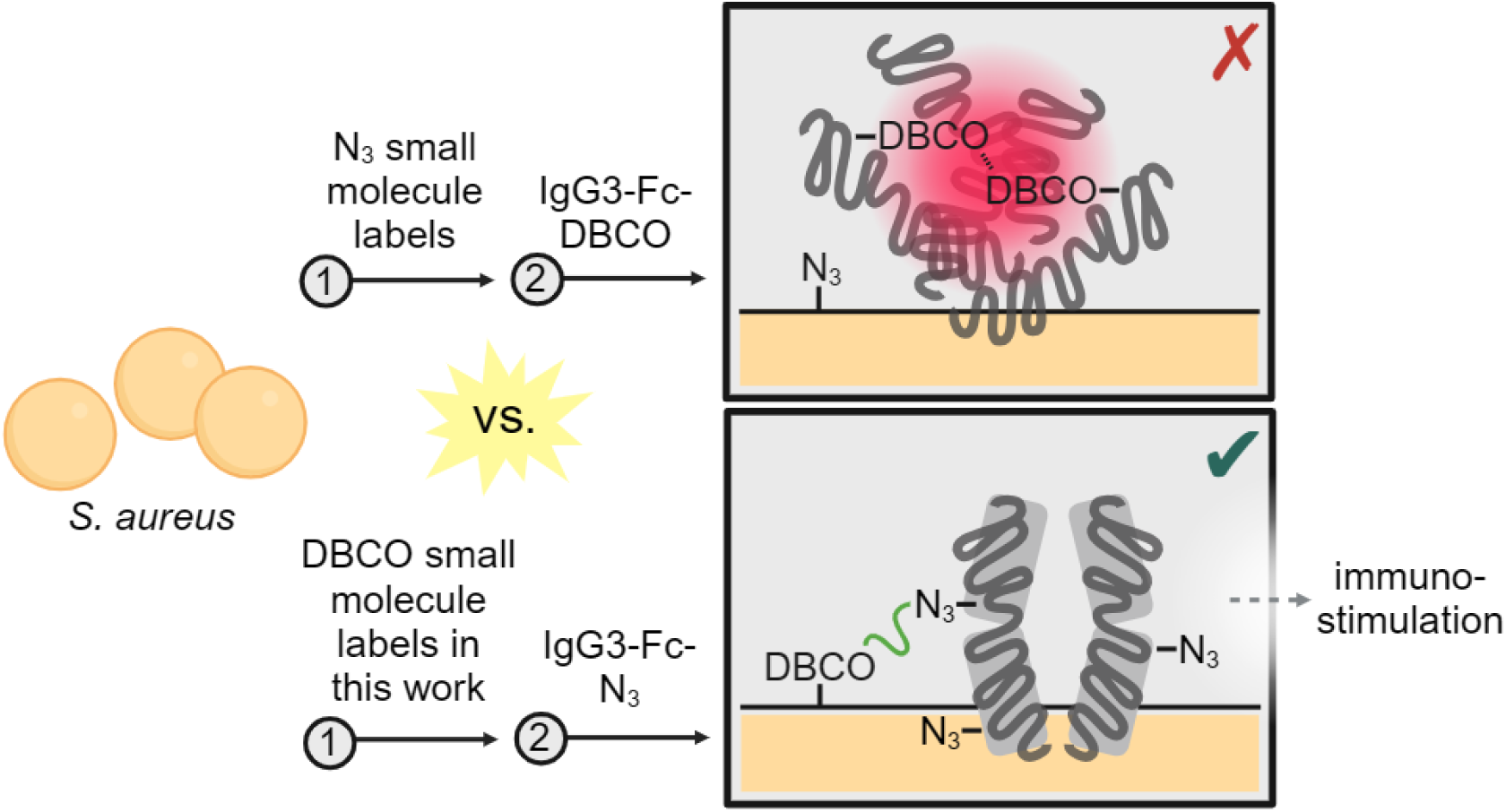

## INTRODUCTION

Click chemistry is a useful tool for modifying biological systems due to its selectivity, efficiency, and bioorthogonality. The copper-free strain promoted azide-alkyne cycloaddition (SPAAC) is a particularly useful cell-compatible reaction and has found widespread use in the fields of cancer drug delivery, transplanted cell tracking, and biomaterial functionalization, among others^1–4^. This reaction is often employed as part of a two-step approach to label living cells that entails installing a small bioorthogonal click group – often, an azide – onto their surface and subsequently delivering a payload (e.g., a protein or nanoparticle) functionalized with the complementary cyclooctyne for *in situ* conjugation. There are several established strategies for introducing an azido group onto the surface of bacteria. One of the most well-validated and widely used strategies, termed metabolic engineering, involves co-incubating bacteria with modified monosaccharides^5–8^, such as the (tetra-acetylated) derivatives *N*-azidoacetylglucosamine (GlcNAz), *N*-azidoacetylmannosamine (ManNAz), and *N*-azidoacetylgalactosamine (GalNAz), the former two being analogs of the conserved disaccharide building blocks of the bacterial peptidoglycan (PGN) and the latter a key component of the polysaccharide capsule in some species. A related but distinct approach is to co-incubate bacteria with modified amino acids, such as azido-D-alanine^5,9–13^, which can be specifically displayed on the surface via either uptake and cytosolic incorporation into PGN precursors and/or transpeptidase-mediated exchange with existing terminal D-alanine residues found in established stem peptide crosslinks.^14^ Labeling bacteria with an accessible azide group and then conjugating cyclooctyne-modified cargo is a strategy that has been used in diverse applications ranging from attaching fluorophores to study PGN biosynthesis and track bacterial growth and migration *in vivo*^5,9^ to attaching heat-generating dyes for ablative photodynamic therapy^10^.

The dibenzocyclooctyne (DBCO) group is one of the most commonly-used cyclooctynes in the SPAAC reaction. Compared to its azide counterpart, the DBCO group is large (molecular weight = 305.3 Da for DBCO-acid vs. 101.1 Da for azidoacetic acid) and hydrophobic (logS = –4.5 for DBCO-NHS vs. –0.2 for azide-NHS), often necessitating that it be dissolved at modestly high concentrations in organic solvents before use, the presence of which can induce destabilization even at low concentrations.^15^ For these reasons, DBCO can cause aggregation issues in sensitive payloads, which can stymie research progress and may go underreported.

These limitations motivated us to explore whether we could label the surface of the model organism *Staphylococcus aureus* with DBCO instead and subsequently co-incubate the bacteria with an azido-labeled protein for easy engraftment. Our payload of interest, human IgG3-Fc, is known to be more prone to aggregation than other subclasses of IgG^16^ due in part to its extended hinge region, and we have empirically observed some degree of aggregation following modification of the protein with DBCO (as discussed later).

Previous efforts to graft related immunostimulants (e.g., IgG proteins or haptens) onto the surface of *Staphylococcus* have relied predominantly on vancomycin as a targeting moiety. Vancomycin is a bactericidal antibiotic that binds the D-alanyl-D-alanine motif in bacterial cell wall precursors with high affinity, preventing further assembly and causing cell death in gram-positive bacteria. For example, Katzenmeyer *et al.* developed a vancomycin-(PLL-g-PEG)-IgG-Fc “artificial opsonin” that led to increased phagocytosis of *Staphylococcus epidermidis* by human neutrophils.^17^ In a similar vein, Sabulski *et al.* created a vancomycin-dinitrophenyl hapten conjugate that also improved phagocyte-mediated killing of *S. aureus*.^18^ Most pertinent to this work, Jiang *et al.* synthesized a DBCO-vancomycin conjugate that selectively bound *S. aureus* over the gram-negative *E. coli* and could subsequently react with an azide-conjugated antibacterial quaternary ammonium compound to induce gram-specific cell death.^19^

Unfortunately, vancomycin has some properties that limit its utility. It is a “last resort” antibiotic that exhibits a strong bacteriostatic effect at moderate concentrations (the minimum inhibitory concentration in *S. aureus* can be from between 0.5 – 2 μg/mL [0.4 – 1.4 μM] in susceptible bacterial populations or upwards of 16 μg/mL [11.2 μM] in resistant ones),^20^ which may not be an acceptable side effect for some surface labeling applications. Moreover, it is ineffective in bacteria with natural or acquired vancomycin resistance, the incidence rate of which has been rising in recent years.^21,22^ For these reasons, we sought to identify alternatives capable of introducing DBCO onto the surface of bacteria that avoid the risks associated with antibiotic use in humans, preserve bacterial viability, and target a conserved and immutable surface pathway to label the PGN of different species.

Sortase A (SrtA) is a transmembrane enzyme found in gram-positive bacteria that catalyzes the covalent anchoring of surface proteins to the cell wall envelope. It recognizes the penta-peptide motif LP*x*TG (where *x* = any amino acid), cleaves between T and G, and transfers the LP*x*T peptide to the PGN.^23^ Incubating of bacteria with LP*x*TG peptides modified at the N-terminus with cargo such as fluorophores, biotin, or click groups (predominantly azides) has been shown to lead to efficient chemical modification of the cell wall in *S. aureus*.^24–27^ The performance of DBCO-modified sortase substrates has not been investigated to the best of our knowledge. As previously mentioned, modified D-amino acids, predominantly D-alanine, are another strategy used to implement small chemical handles into the PGN of bacteria, although the performance of DBCO-modified D-alanine has similarly not been reported.

We investigated the effectiveness of these two types of DBCO-modified small molecules – a sortase substrate peptide and D-alanine – in labeling the surface of *S. aureus* with both a fluorophore (sCy5-azide) and IgG3-Fc protein labeled with a fluorophore (IgG3-Fc-sCy5-azide) and compared their incorporation efficiency to that of the established DBCO-vancomycin conjugate. Additionally, we investigated off-target labeling and toxicity in mammalian cells to gauge the translational relevance of these approaches.

## RESULTS AND DISCUSSION

### Synthesis of DBCO-Modified Small Molecules

We synthesized the DBCO-modified sortase substrate (LPETG, **1**) and scramble control (EGTLP, **2**) via standard Fmoc solid phase peptide synthesis (SPPS) on rink amide MBHA resin, using (MeCN)_4_CuBF_4_ to protect the DBCO group from acid-catalyzed rearrangement during cleavage^28^ (see Materials and Methods). Note that it has also been demonstrated that the substrate sequence LPMTG (with FITC at the N-terminus) can achieve a higher degree of labeling in *S. aureus*.^27^ However, the strategy we used to protect the DBCO group from acid-mediated rearrangement during the cleavage step of SPPS involves using (MeCN)_4_CuBF_4_, a copper-containing compound. Methionine (M) is the only canonical amino acid that is not fully compatible with these conditions as it is susceptible to copper-catalyzed oxidation and can cause the formation of undesirable side products. To circumvent this problem, we opted to use the canonical LPETG sequence and corresponding scramble control, EGTLP, instead of the LPMTG/MGTLP pair. We synthesized the DBCO-modified L- and D-alanine precursors (DBCO-Boc-Dap-OH and DBCO-Boc-D-Dap-OH, respectively) using NHS ester-amine chemistry. In addition to testing a variant with a conventional carboxylic acid (COOH) group at the C-terminus, we also sought to test the effect of including an amide group (CONH_2_) instead, as it has previously been shown that this modification can improve the incorporation of modified D-amino acids into the bacterial PGN.^29^ Thus, the precursors were subsequently loaded onto rink amide MBHA resin or left as-is and treated with a cleavage/deprotecting cocktail to generate the amide-terminated variants **3a** and **4a** or carboxylic acid-terminated variants **3b** and **4b**, respectively. Finally, the DBCO-vancomycin conjugate **5** was synthesized via NHS-amine chemistry. All synthesized compounds (summarized in **Figure 1**) were purified via reversed-phase high-performance liquid chromatography (RP-HPLC). To confirm that the acid-sensitive DBCO group remained intact during synthesis, a sample of each compound was reacted with an excess of 6-azido-hexanoic acid to demonstrate a shift in the reversed-phase ultra high-performance liquid chromatography (RP-UPLC) peak at 220/230 nm and complete ablation of the peak at 307 nm, signifying the formation of the DBCO-azide complex (**Figure S1**).

**Figure 1.**
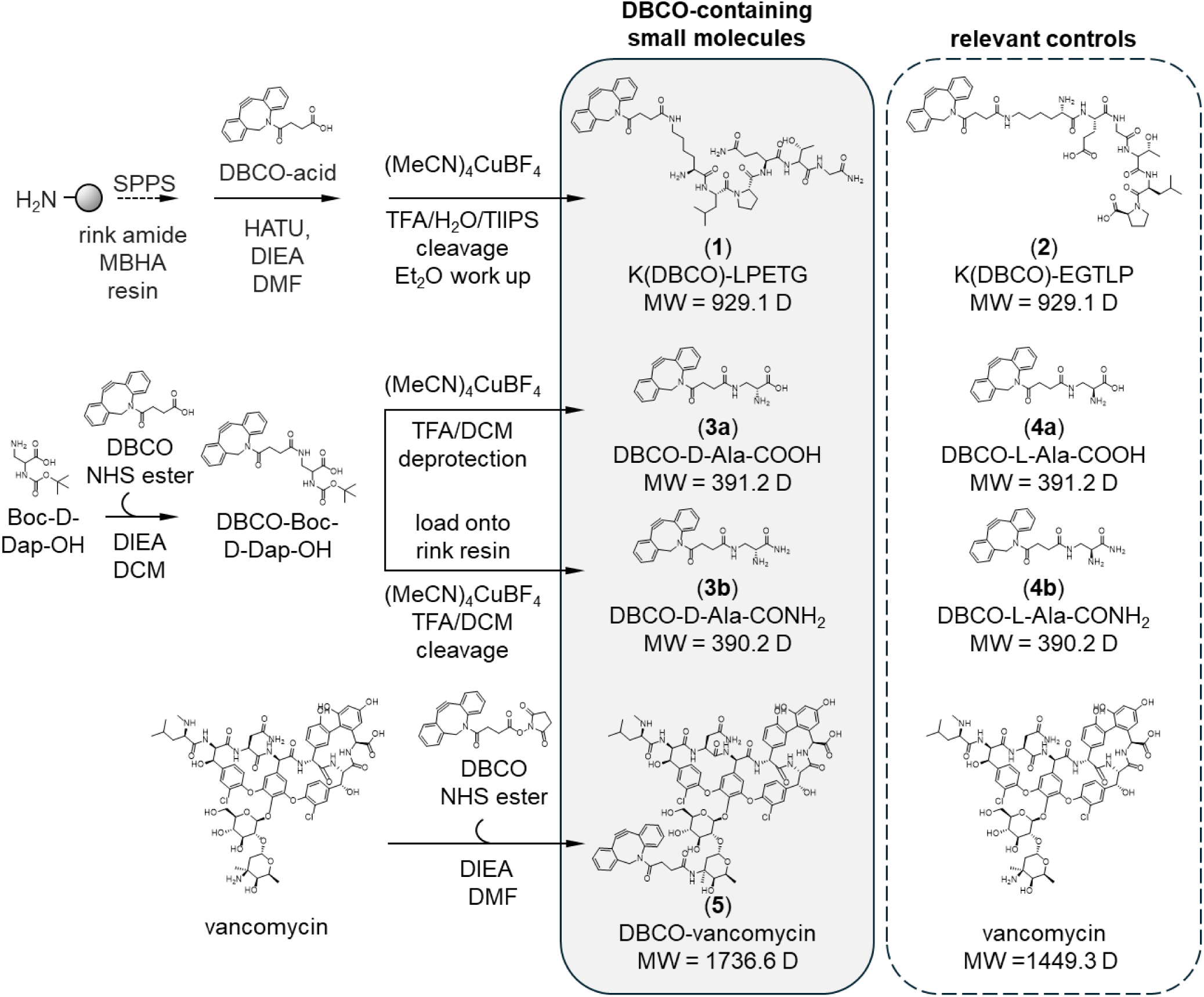
Synthesis schemes for the small molecules used in this work. Compounds **1** and **2** were synthesized via standard Fmoc solid-phase peptide synthesis on rink amide MBHA resin. Compounds **3a**, **4a**, **3b**, and **4b** were synthesized via an NHS-amine reaction between Boc-(D)-Dap-OH and DBCO-NHS, followed optionally by loading onto rink amide MBHA resin to generate the amide terminated variants (**3b** and **4b**). All five compounds were then treated with (MeCN)_4_CuBF_4_ to prevent acid-catalyzed rearrangement of the DBCO group prior to cleavage off of resin and/or deprotection of the Boc group with 30:70 solution of TFA:DCM. DBCO-vancomycin (**5**) was similarly synthesized using NHS-amine chemistry, sans a cleavage or deprotection step. All compounds were purified using RP-HPLC and characterized via RP-UPLC and MS-ESI prior to use.

### Determining labeling efficiency in *S. aureus*

We proceeded to evaluate the ability of each small molecule to label the surface of *S. aureus* Newman. *S. aureus* was co-incubated with 50 μM of each small molecule for 24 hours. For the DBCO-vancomycin (**5**) and vancomycin-only groups, the starting OD600 of the cultures was set to 1 instead of 0.1, as designated^†^, in order to counteract the bacteriostatic effects of vancomycin so that the total number of cells was comparable between samples at the end of the incubation period. Bacteria were then fixed with 2% paraformaldehyde (PFA) in phosphate-buffered saline (PBS) and resuspended in a solution containing 1 μM sulfo Cy5-azide (sCy5-azide) for 2 hours at room temperature (RT) and analyzed via flow cytometry (**Figure 2**). The DBCO-modified sortase substrate **1** was initially unable to specifically label the bacteria over the scramble control **2**. Both compounds were additionally tested at a higher concentration of 500 μM, which did produce a significant difference (geometric mean fluorescent intensity [MFI] = 145 ± 2 AU for 500 μM of **1** vs. 40 ± 2 AU for 500 μM of **2**). This is not unexpected in hindsight, given that 0.25 – 1 mM of sortase substrate is commonly used in similar assays.^24–27^ Therefore, this approach was reasonably specific, but only efficient at modestly high concentrations. Both compounds **3a** and **4a**, the carboxylic acid-terminated DBCO-D-alanine and DBCO-L-alanine, respectively, were able to label the bacteria (MFI = 1,108 ± 56 AU for **3a** vs. 307 ± 11 AU for **4a**), with an evident stereoselective advantage for the D-enantiomer over the L-enantiomer. Compound **3b**, the amine-terminated DBCO-D-alanine, produced an increase in fluorescence (MFI = 189 ± 117 AU) but was less efficient compared to its carboxylic acid-terminated counterpart. Interestingly, its stereocontrol, **4b**, produced little-to-no labeling (MFI = 21 ± 2 AU), suggesting an improvement in stereoselectivity at the expense of absolute labeling efficiency. The labeling ability of compounds **3a** and **3b** was surprising given the comparatively large size and hydrophobic nature of the DBCO group. In line with previous findings, DBCO-vancomycin was able to strongly label the surface of the *S. aureus* (MFI = 36,232 ± 848 AU) whereas vancomycin itself had no statistically significant effect (MFI = 22 ± 2), confirming that the interaction between vancomycin and the PGN itself did not increase the bacteria’s susceptibility to non-specific labeling by the sCy5-azide. Thus, compounds **1**, **3a**, **4a**, **4b**, and **5** were all able to label *S. aureus* to different extents (**5** >> **3a** > **4a** ⪎ **3b** >> **4b** ≈ 0) and, of the DBCO-L/D-alanine variants, the carboxylic acid-terminated compounds outperformed their amide-terminated counterparts in terms of absolute labeling efficiency but exhibited less stereospecificity for D-vs. L-alanine. The vancomycin-DBCO achieved the highest labeling efficiency of the tested compounds when the starting OD600 of the culture was preemptively increased ten-fold.

**Figure 2.**
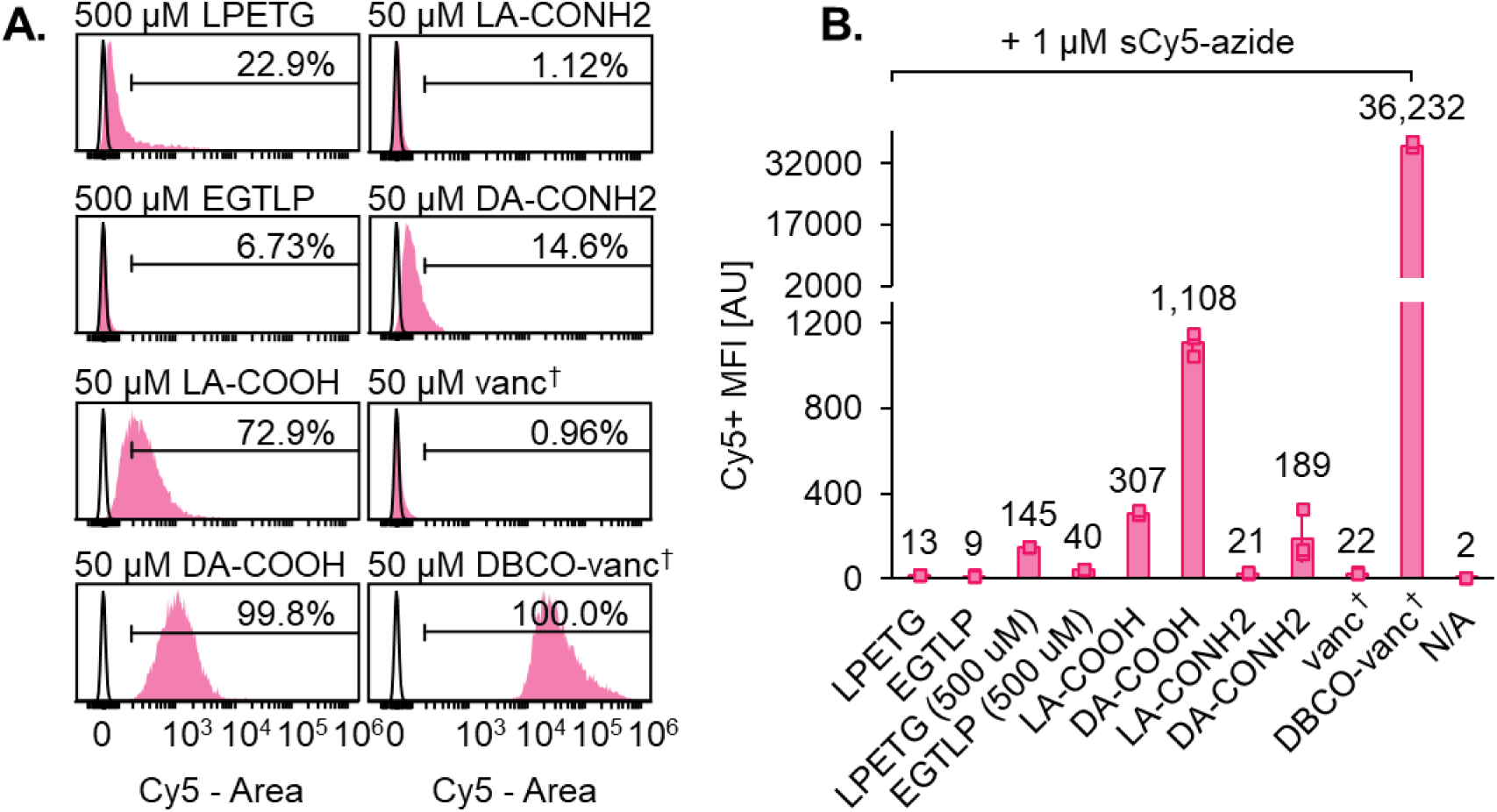
Labeling efficiency in *S. aureus* using DBCO-containing small molecules. *S. aureus* concentration was normalized to a starting OD600 of 0.1 or 1.0 (as signified^†^, to counteract the bactericidal effects of DBCO-vancomycin and vancomycin) and co-incubated with 50 or 500 μM of the specified DBCO-containing small molecules for 24 hours at 37 °C. Afterwards, the bacteria was fixed and co-incubated with 1 μM sCy5-azide for 2 hours at room temperature prior to analysis via flow cytometry. The histograms to the left (**A**) are representative samples from the data set summarized in (**B**), plotted as the geometric mean fluorescent intensity (MFI) of three experimental replicates. Compounds **1**, **3a**, **4a**, **3b**, and **5** were all able to label *S. aureus* to different extents (**5** >> **3a** > **4a** > **3b** >> **4b** ≈ 0).

### Determining effect on bacterial growth

We next sought to investigate the effects of these compounds on the viability and growth of *S. aureus* Newman. We co-incubated the bacteria with 50 and/or 500 μM of compounds **1**, **2**, **3a**, **4a**, **3b**, **4b**, **5**, or vancomycin, and periodically measured the OD600 as an indicator of cell growth (**Figure 3**). As expected, DBCO-vancomycin (**5**) and vancomycin had a stark bacteriostatic effect (OD600 = 0.121 ± 0.001 for **5** and 0.125 ± 0.002 for vancomycin compared to 0.51 ± 0.03 for the vehicle control after 12 hours). In contrast, the remaining compounds did not impact bacterial growth or caused a small but statistically significant increase in OD600 after 12 hours of incubation in the case of 500 μM of compound **1** (OD600 = 0.60 ± 0.01) and both 50 and 500 μM of compound **2** (OD600 = 0.56 ± 0.01 and 0.58 ± 0.01, respectively). These data show that a subset of compounds (**1**, **3a**, **3b**, and **4a**) can label the surface of *S. aureus* with accessible DBCO without affecting the growth of the bacteria.

**Figure 3.**
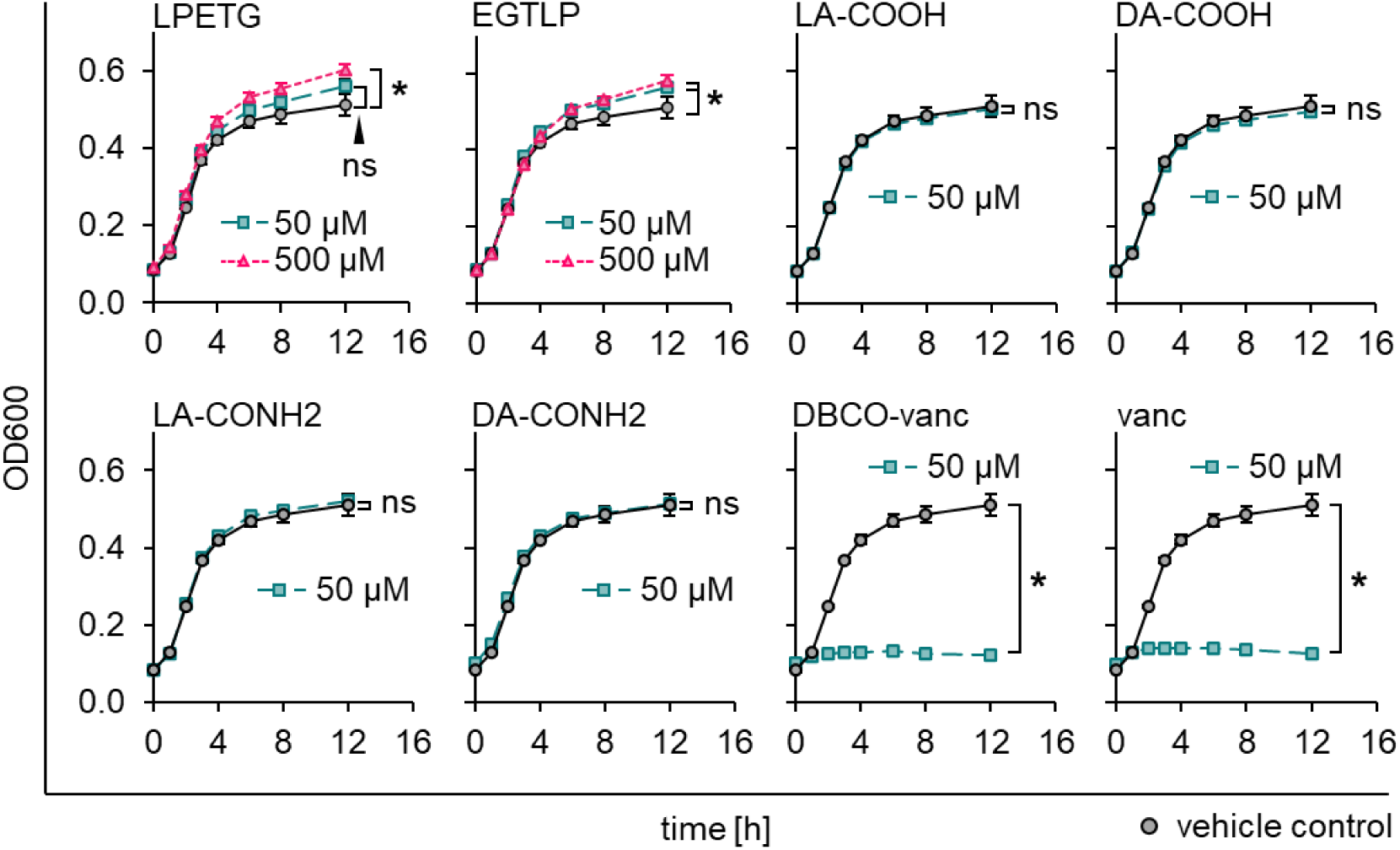
Effect of DBCO-containing small molecules on the growth rate of *S. aureus*. *S. aureus* was normalized to a starting OD600 of 0.1 and co-incubated with 50 or 500 μM of the specified DBCO-containing small molecule in a plate reader for 12 hours at 37 °C. The OD600 was measured every hour as a proxy for bacterial growth. The novel compounds (**1**, **2**, **3a**, **3b**, **4a**, and **4b**) had no effect or a slightly beneficial effect on bacterial growth whereas DBCO-vancomycin (**5**) and vancomycin were starkly bacteriostatic.

### Determining off-target labeling and toxicity at high concentrations in HEK293T cells

Given that many *in situ* cell labeling applications require selectively targeting bacteria in a complex environment that additionally contains mammalian cells, we next tested the bacterial specificity of the DBCO-modified small molecules. We excluded the DBCO-modified sortase substrate **1** (and its stereocontrol **2**) from further testing due to its poor labeling efficiency compared to compounds **3a** and **3b** with no apparent advantage for the specific application at hand (grafting proteins onto the surface of a single species of bacteria *in vitro*). This approach may remain useful in applications where species selectivity is desired, as *S. aureus* is the archetypal bacteria for studying SpA expression.^30^ We co-incubated human embryonic kidney HEK293T cells with 50 or 500 μM of each molecule for 16 hours, then stained the cells with 1 μM sCy5-azide (**Figure 4A**). Of the four DBCO-modified L/D-alanine compounds tested, compound **4b** was the only one that led to a statistically significant increase in off-target labeling at the higher concentration (500 μM), with 27 ± 7% of HEK293T singlets gated as Cy5+, compared to 8 ± 2% for the vehicle control (*p* = 0.01). This compound was incidentally unable to label *S. aureus* (at 50 μM). DBCO-vancomycin (**5**) also led to off-target labeling at the higher concentration tested, with 56 ± 6% of singlets gated as Cy5+ (*p* = 0.001), which could be problematic for applications involving mixed cell populations where bacterial specificity is key, especially in *in vitro* or *ex vivo* cultures, where higher concentrations of DBCO-vancomycin may be more feasible/better tolerated. We also performed a colorimetric 3-(4,5-dimethylthiazol-2-yl)-5-(3-carboxymethoxyphenyl)-2-(4-sulfophenyl)-2H-tetrazolium (MTS) assay to quantify the metabolic activity of the HEK293T cells after treatment with compounds **3a**, **4a**, **3b**, **4b**, and **5** as a proxy for cell viability (**Figure 4B**). In line with the results of the co-incubation experiment, compounds **4b** and **5** both caused significant decreases in metabolic activity, potentially due to non-canonical amino acid integration into proteins and subsequent interference with cellular processes for the former. One caveat, however, is that the conditions studied here are higher than physiologically-relevant concentrations of vancomycin-DBCO or vancomycin, the latter of which has been used in the range of 3 – 30 μM;^31^ therefore, these levels would not be expected to cause significant cellular toxicity and our results therefore constitute an edge case. In summary, compounds **3a**, **4a**, and **3b**, all of which are capable of labeling bacteria with accessible DBCO moieties without affecting bacterial growth, did not lead to statistically significant labeling or cellular toxicity in mammalian cells at high concentrations.

**Figure 4.**
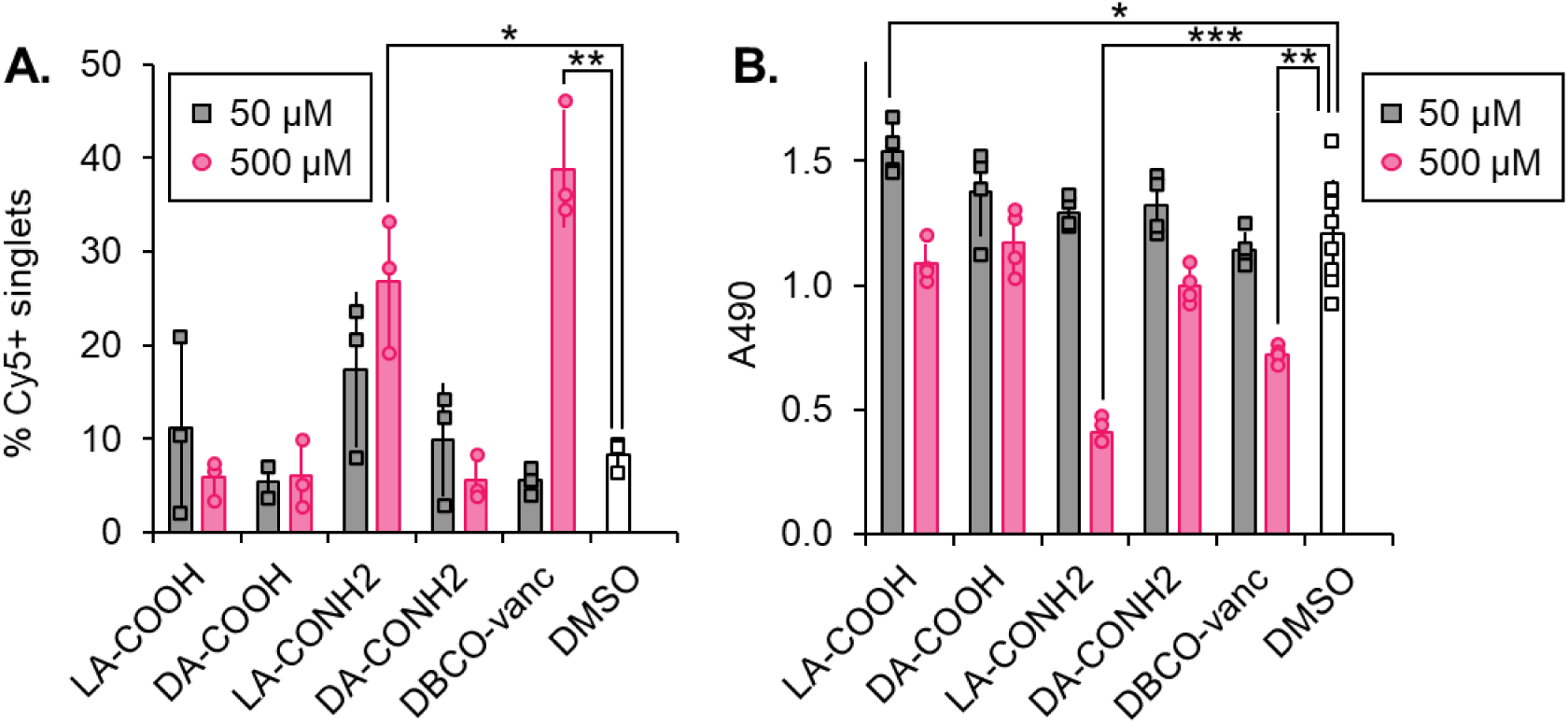
Off-target labeling and toxicity in mammalian cells of DBCO-containing small molecules. HEK293T cells were co-incubated with 50 or 500 μM of the specified DBCO-containing small molecules for 16 hours at 37 °C. Afterwards, the cells were (**A**) co-incubated with 1 μM sCy5-azide for 30 minutes at 4 °C and analyzed via flow cytometry or (**B**) co-incubated with MTS reagent for 1 hour at 37 °C and the absorbance of each well at 490 nm was measured using a plate reader. Compounds **3a**, **4a**, and **3b** did not lead to statistically significant labeling or cause cellular toxicity in mammalian cells at either concentration whereas compounds **4b** and **5** both labeled the cells and caused a significant decrease in metabolic activity at the higher of the two concentrations tested.

### Labeling IgG3-Fc with azido groups and assessing protein conjugate characteristics

We next sought to leverage the DBCO handle introduced into the PGN of *S. aureus* to install a protein of interest onto the cell surface. We chose IgG3-Fc for its therapeutic relevance. Staphylococcal protein A (SpA) present on the surface of *S. aureus* can bind the constant region (Fc) of human IgG1, IgG2, and IgG4, misorienting them. This misorientation prevents them from properly binding Fc receptors on the surface of immune cells, stymieing antibody- and complement-dependent cellular cytotoxicity. However, SpA cannot bind human IgG3 due to the H435R mutation in the Fc region; accordingly, circulating IgG3 is known to play an important role in humoral immunity against *S. aureus*.^32^ Using our approach, we sought to attach azide-modified IgG3-Fc to the surface of bacteria, creating a ‘pseudo-antibody’ with specificity dictated by pre-facto labeling of the bacterial PGN with DBCO.

In earlier experiments, we empirically observed that modifying IgG3-Fc with DBCO led to issues with protein aggregation. To illustrate this, we labeled IgG3-Fc with 6 molar equivalents (eq.) of sulfo DBCO-PEG_4_-TFP (sDBCO-PEG_4_-TFP), 6 eq. of azide-PEG_4_-NHS, or DMSO only (vehicle control) and stored the protein samples at 4 °C, RT, or 37 °C for 72 hours. When ran the samples on an SDS-PAGE protein gel and observed that the unlabeled protein sample and the sample labeled with azide produced a single band when stored at 4 °C or RT but two bands (one having a higher MW) when both samples were stored at 37 °C, comprising 2 – 3% of the total protein content in either case. In comparison, the protein sample labeled with DBCO produced a second band at all three storage temperatures, comprising 5 – 9% of the total protein content (**Figure S2**). This apparent aggregation could lead to non-specific protein adhesion, which could have undesirable effects when the intended application is, for example, directing the deposition of an immunostimulatory molecule.

We proceeded to use the related small molecule, sCy5-azide-NHS, to label the IgG3-Fc with an accessible azide moiety. Use of this bifunctional and fluorescent molecule allowed us to quickly obtain an estimate of the degree of labeling (DOL) by measuring the absorbance of the protein sample at 280 and 647 nm after the removal of unbound material and easily detect IgG3-Fc deposited on the surface of bacteria using flow cytometry without the need to use a secondary antibody that would be subject to the aforementioned criteria to avoid nonspecific surface adhesion by SpA. As with the azide-PEG_4_-NHS, the sCy5-azide-NHS molecule did not cause aggregation issues at any of the concentrations tested. We ultimately chose to label IgG3-Fc with 3 eq. (“low” labeling) and 4 eq. (“high” labeling) of Cy5-azide-NHS; these equivalencies were chosen after determination of the DOL such that ≥ 80% and ≥ 95% of IgG3-Fc molecules were labeled with at least one sCy5-azide molecule, respectively.

For a high-fidelity estimate of the DOL, we reacted the modified IgG3-Fc-sCy5-azide protein (or unmodified IgG3-Fc protein) with an excess of 5 kDa mPEG-DBCO and ran an SDS-PAGE protein gel (**Figure 5A**). The IgG3-Fc-sCy5-azide protein produced a ladder-like band pattern, with consecutive bands corresponding to the addition of zero, one, two, three, etc. DBCO-mPEG groups. Thee bands were separated more than their standalone molecular weights would suggest since the PEG groups form complexes with SDS, restricting their travel through the gel.^33,34^ We quantified the intensity of each adduct band and fit a Poisson curve to the resulting distribution, setting the λ parameter equal to the DOL. Using this method, we calculated DOL values of 1.7 and 3.9 for the so-called “low” and “high” labeling groups, respectively (**Figure 5B**).

**Figure 5.**
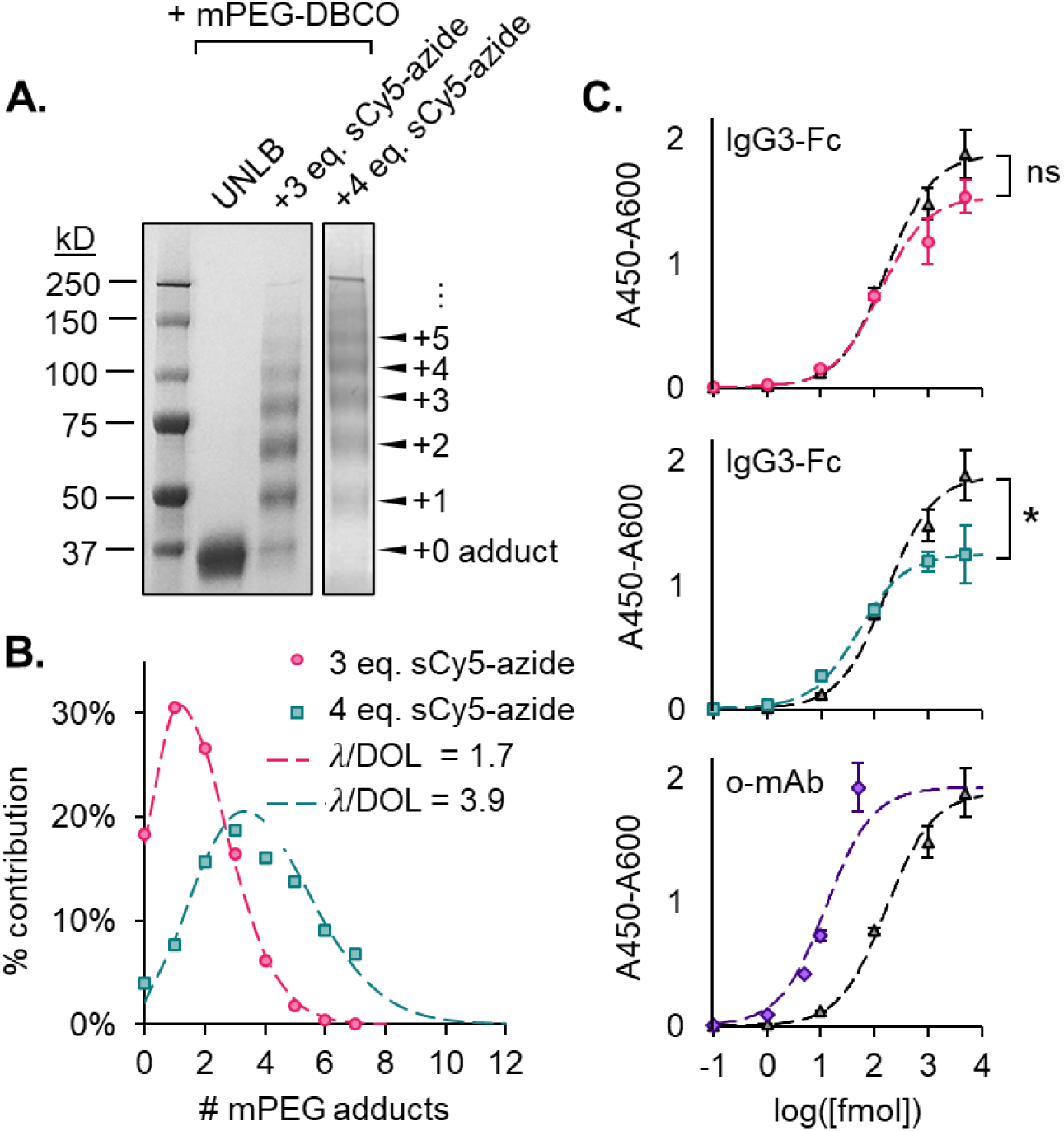
Labeling of IgG3-Fc with azido groups and assessing protein conjugate characteristics. (**A**) Human IgG3-Fc was co-incubated with 3 or 4 molar equivalents (eq.) of sCy5-azide-NHS for 2 hours at RT, then passed through a size-exclusion spin column to remove unbound material. An aliquot of each protein was then reacted with a large excess of 5 kDa mPEG-DBCO and run on an SDS-PAGE gel to determine the degree of labeling (DOL). (**B**) The intensity of each adduct band in (**A**) was quantified via ImageJ and a Poisson curve was fit to the resulting data distribution to yield the λ parameter, which is synonymous with the DOL. (**C**) A sandwich ELISA was performed to determine whether the sCy5-azide-labeled IgG3-Fc and sCy5-labeled Omodenbamab (o-mAb), an anti-SpA IgG3 antibody, retained their ability to bind CD64. IgG3-Fc-sCy5-azide did not exhibit a significant difference in binding to CD64 at a DOL of 1.7 compared to unlabeled IgG3-Fc and exhibited a small but significant reduction in binding at a DOL of 3.9. This suggests that IgG3-Fc labeled with a moderate amount of sCy5-azide retains the ability to perform one of its key effector functions, resulting in activation of an immune response.

Next, we sought to confirm that the addition of the sCy5-azide groups did not disrupt the IgG3-Fc protein’s ability to bind human CD64 (a.k.a. FcγRI), which can trigger an immune response. CD64 is found on the surface of macrophages and activated neutrophils and binds to monomeric IgG with high affinity, serving as a blood marker for various inflammatory disease states including bacterial infections^35^ and sepsis.^36^ We also tested the binding of full-length omodenbamab (o-mAb), a humanized anti-SpA IgG3 antibody in clinical development, labeled using 6 eq. sulfo Cy5-NHS (sCy5-NHS) (estimated DOL = 1.9 via 647 and 280 nm absorbance measurements). As shown in **Figure 5C**, an enzyme-linked immunosorbent assay (ELISA) demonstrated that IgG3-Fc-sCy5-azide did not exhibit significant differences in binding to CD64 at the lower DOL (1.7) compared to unlabeled IgG3-Fc and exhibited a small but significant reduction in binding at the higher DOL (3.9). This trend – an increase in sCy5-azide DOL resulting in a commensurate decrease in CD64 binding – appeared to remain consistent for higher DOLs tested (**Figure S3**). The ELISA also demonstrated that sCy5-labeled o-mAb strongly bound CD64, as anticipated.

### Performing two-step labeling to graft IgG3-Fc onto the surface of *S. aureus*

Finally, we tested whether functional IgG3-Fc could be deposited onto the surface of *S. aureus* by installing DBCO groups on the bacterial surface and subsequently co-incubating the bacteria with IgG3-Fc-sCy5-azide (DOL = 1.7). *S aureus* was co-incubated with 50 μM of compounds **3a**, **4a**, **3b**, **4b**, or **5**, or with 1 μM Cy5-labeled o-mAb for 24 hours at 37 °C, then fixed with 2% PFA in PBS and additionally co-incubated with or without 1 μM IgG3-Fc-sCy5-azide (DOL = 1.7) for 2 hours at RT. As shown in **Figure 6**, there was a significant increase in fluorescence for compounds **3a**, **4a**, **3b**, **5**, and the sCy5-labeled o-mAb. The absolute increase in fluorescence intensity was lower when IgG3-Fc-sCy5-azide was used as the secondary label as compared to sCy5-azide by a consistent factor of approximately 30 – 35, ostensibly due to the fact that a comparatively bulky IgG3-Fc protein (MW ∼ 31,200 Da) labeled with, on average, 0 – 3 molecules of sCy5-azide (MW ∼ 840 Da) obstructs a swath of PGN that could otherwise be occupied by dozens of standalone sCy5-azide molecules. However, the relative apparent labeling efficiencies (**5** >> **3a** > **3b** ⪎ **4a >> 4b** ≈ 0) remained the same regardless of the secondary label used, further corroborating these results.

**Figure 6.**
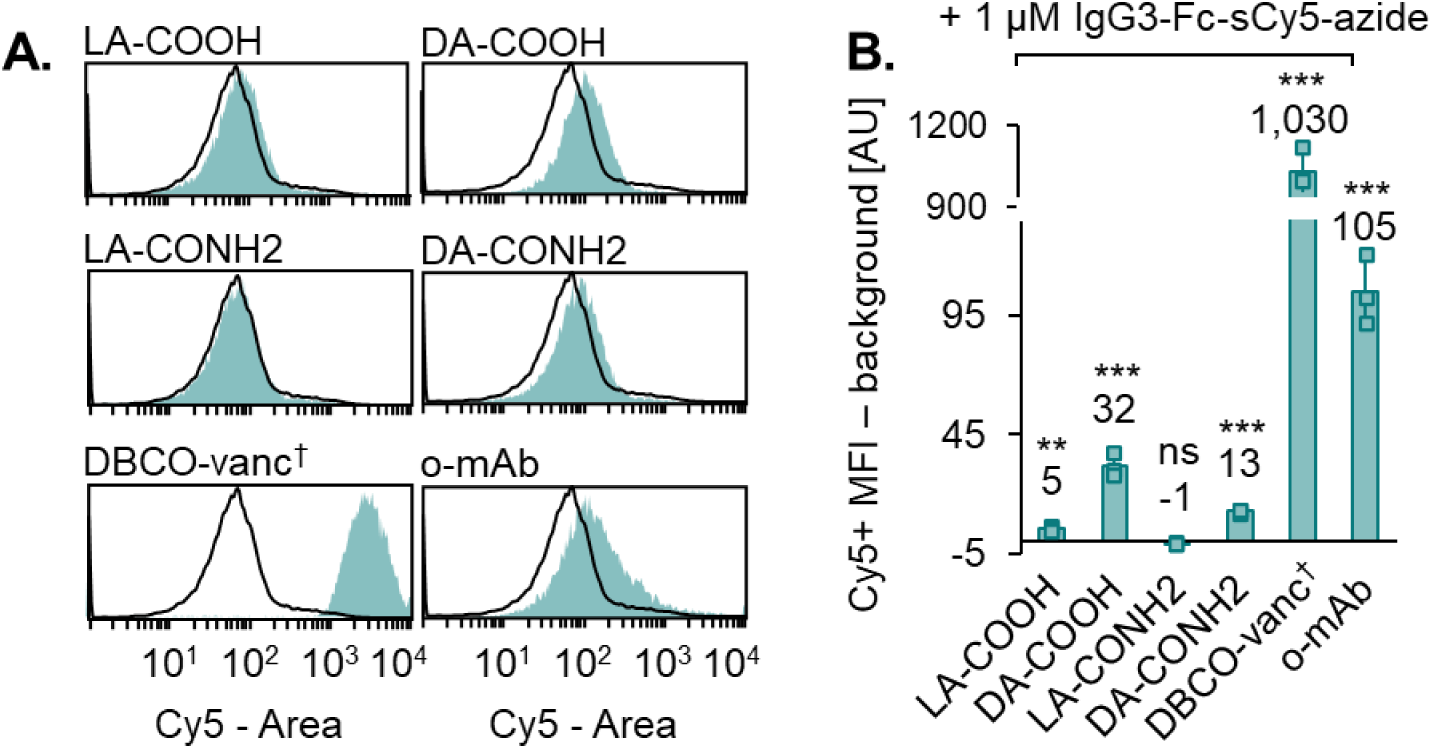
Grafting IgG3-Fc-sCy5-azide onto the surface of *S. aureus*. *S. aureus* was normalized to a starting OD600 of 0.1 or 1.0 (as signified^†^, to counteract the bactericidal effects of DBCO-vancomycin) and co-incubated with 50 μM of the specified DBCO-containing small molecules or 1 μM sCy5-labeled Omodenbamab (o-mAb) aerobically for 24 hours at 37 °C. Afterwards, the bacteria was fixed and optionally co-incubated with 1 μM IgG3-Fc-sCy5-azide (DOL = 1.7) for 2 hours at room temperature prior to analysis via flow cytometry. The histograms to the left (**A**) are representative samples from the data set in (**B**), plotted as the geometric mean fluorescent intensity (MFI) of three replicates with \**p* < 0.05, \*\**p* < 0.01, \*\*\**p* < 0.001 compared to the DBCO-untreated control (co-incubated with 1 μM IgG3-Fc-sCy5-azide only). The trends in relative apparent labeling efficiencies for the small molecule DBCO labels (**5** >> **3a** > **3b** ⪎ **4a** >> **4b** ≈ 0) are the same as those observed when using standalone sCy5-azide.

## CONCLUSIONS

Taken together, our results show that DBCO can be successfully incorporated into the PGN of *S. aureus* using a DBCO-modified sortase substrate and D-alanine amino acid (terminated with either a carboxylic acid or amide) and validate prior work using a DBCO-vancomycin conjugate. Of the novel small molecule labels explored in this work, the carboxylic acid-terminated D-alanine (**3a**) led to the highest degree of labeling that was several times greater than that of the L-enantiomer control (**4a**). In comparison, the amide-terminated D-alanine variant (**3b**) led to less labeling but improved stereoselective incorporation, such that the L-enantiomer control (**4b**) was effectively inert in bacteria. Compared to the performance of DBCO-vancomycin (**5**), the DBCO-modified D-alanine variants **3a**, **4a**, and **3b** did not perturb the growth of the *S. aureus* and did not cause any off-target labeling or toxicity in mammalian cells, even at high concentrations. Conversely, compound **4b** led to a comparatively high degree of labeling in mammalian cells while causing a commensurate decrease in cell metabolic activity/viability, suggesting the potential to selectively target mammalian cells over bacteria.

We were able to label human IgG3-Fc protein with sCy5-azide as well as azide-PEG_4_. Unlike IgG3-Fc labeled with sDBCO-PEG_4_, IgG3-Fc labeled with sCy5-azide/azide-PEG_4_ did not meaningfully aggregate under normal storage conditions. Additionally, IgG3-Fc-sCy5-azide (DOL = 1.7) retained its ability to bind CD64. We were able to use this molecule to install functional IgG3-Fc onto the surface of *S. aureus* treated with compounds **3a**, **4a**, **3b**, and **5**.

We believe the labeling strategies explored herein will prove useful for modifying the surface of *S. aureus* and other gram-positive bacteria via the SPAAC reaction when the proteins and other cargo of interest are prone to destabilization or otherwise incompatible with DBCO modification. Examples include proteins recombinantly synthesized to contain an azidonorleucine^37,38^ (e.g., antibodies or antigen-binding ligands) or peptide-based supramolecular nanostructures encapsulating a drug that can accommodate an azide moiety but not a cyclooctyne without experiencing substantial disruption to their secondary structure. The labeling strategies involving DBCO-modified D-alanine may be more suitable over vancomycin-DBCO when the application at hand necessitates that the bacteria remain metabolically active, such as when harnessing innate chemosensory pathways to direct engineered cells to a site of interest (and/or produce a therapeutic payload) or when monitoring commensals without intending to induce dysbiosis. Accordingly, using DBCO-vancomycin may be more appropriate when it is desirable to achieve extremely high levels of surface labeling and bacterial viability is not important (for example, if using dead/replication-incompetent whole pathogenic bacteria or PGN fragments conjugated to an antigen for vaccine applications). Finally, our results provide insight into the design rules for creating modified sortase substrates and D-amino acids for labeling applications and, given the effectiveness of both DBCO-D-alanine variants, demonstrate some unexpected permissibility in the latter approach.

## EXPERIMENTAL PROCEDURES

### Synthesis of DBCO-LPETG/EGTLP sortase substrates

DBCO-modified sortase peptide substrates **1** and **2** were synthesized via standard Fmoc solid phase peptide synthesis (SPPS) using hexafluorophosphate azabenzotriazole tetramethyl uronium (HATU)/ N,N-diisopropylethylamine (DIEA) coupling chemistry on rink amide MBHA resin. The resin was deprotected using 25 v/v% piperidine in N,N-dimethylformamide (DMF). Four molar equivalents (eq.) of each Fmoc-protected amino acid, 4 eq. HATU, and 6 eq. DIEA dissolved in a minimal amount of DMF were added to the resin and reacted for 20 minutes. Afterwards, the resin was washed with dichloromethane (DCM) and DMF to remove any unreacted material. This cycle of deprotection-coupling was repeated for the remaining five amino acids, with the Boc-Lys(Fmoc)-OH spacer added last. Following deprotection, 2.5 eq. DBCO-acid along with 2.5 eq. HATU and 3.75 eq. DIEA in DMF was added to the resin for 1 hour. To protect the DBCO group from acid-catalyzed rearrangement during the subsequent cleavage step, 2 eq. (MeCN)_4_CuBF_4_ was added as a dry powder to the resin immediately prior to the addition of 95:2.5:2.5 TFA:H_2_O:triisopropylsilane for 2 hours. Most of the cleavage cocktail was subsequently evaporated off and the crude material was precipitated in cold diethyl ether. The solution was centrifuged at 3,000 relative centrifugal force (RCF) for 5 minutes at 4 °C, the supernatant was decanted, and the pelleted material was air-dried at RT. The crude material was purified as described below.

### Synthesis of DBCO-L/D-alanine-(COOH/CONH_2_)

DBCO-NHS (20 mg, 1 eq.) and either Boc-Dap-OH or Boc-D-Dap-OH (10.2 mg, 1 eq.) were combined in 2 – 3 mL of DCM with DIEA (26 μL, 3 eq.). The reactions were stirred at RT overnight and then split in half, yielding a total of four aliquots, two containing DBCO-Boc-Dap-OH and two containing DBCO-Boc-D-Dap-OH. The DCM and DIEA were evaporated off using a solvent evaporator. To half of the aliquots, (MeCN)_4_CuBF_4_ (15.7 mg, 2 eq.) was added as a dry powder and the material was resuspended in 2 mL of a 30:70 solution of TFA:DCM and allowed to stir at RT for 2 hours to remove the Boc protecting groups, generating the carboxylic acid-terminated compounds **3a** and **4a**. To the other half of the aliquots, HATU (28.4 mg, 3 eq.) and DIEA (26 μL, 6 eq.) were added and the material was resuspended in 2 – 3 mL of DMF. The solution was then added to deprotected rink amide MBHA resin and allowed to couple for 1 hour at RT. Afterwards, (MeCN)_4_CuBF_4_ (15.7 mg, 2 eq.) was added as a dry powder to the resin and the compounds were deprotected/cleaved with a 30:70 solution of TFA:DCM for 2 hours. Most of the TFA and DCM were evaporated off using a solvent evaporator and the crude material was precipitated in cold diethyl ether. The solution was spun at 3,000 RCF for 5 minutes at 4 °C, the supernatant was decanted, and the pelleted material was air-dried at RT. The crude material was purified as described below.

### Synthesis of vancomycin-DBCO

Vancomycin-HCl (30 mg, 1 eq.) and DBCO-NHS (8.5 mg, 1.05 eq.) were combined in 5 – 10 mL of DMF with DIEA (21.2 μL, 6 eq.) and the mixture was stirred at RT for 24 hours. Afterwards, the DMF and DIEA were completely evaporated off using a solvent evaporator. The crude material was purified as described below.

### General purification and characterization of DBCO-containing small molecules

Crude products were resuspended in 50:50 acetonitrile (ACN):H_2_O and purified via RP-HPLC using a Shimadzu C18 HPLC column with H_2_O with 0.05% trifluoracetic acid (TFA) (A)/ACN with 0.05% TFA (B) as the solvent system. The method used was a 20 – 70%B gradient with a 3%B/minute ramp rate. The ACN was then completely evaporated off using a solvent evaporator and the solution was sterile-filtered and lyophilized. The pure materials were dissolved in DMSO at a stock concentration of 30 – 100 mM.

The purified products were characterized via RP-UPLC using a 1290 Infinity II instrument (Agilent, Santa Clara, California, USA) with a Poroshell 120 EC-C18 column and the aforementioned solvent system. The method used was a 5 – 95%B gradient with a 15%B/minute ramp rate. The products were also characterized via mass spectrometry electrospray ionization (MS-ESI) using an InfinityLab LC/MSD iQ single quadrupole instrument (Agilent, Santa Clara, California, USA).

To confirm the reactivity of the DBCO group, a small amount of each material (∼0.1 μg) was reacted with a large excess of 6-azido hexanoic acid (∼2 μg) in a small volume (∼100 – 200 μL) overnight at RT and analyzed via RP-UPLC (**Figure S1**).

### Co-incubating *S. aureus* with DBCO-containing small molecules (primary labels)

*S. aureus* Newman was streaked onto a tryptic soy agar (TSA) plate and grown overnight (16 – 20 hours) at 37 °C. A single colony was used to inoculate a 10 mL liquid tryptic soy broth (TSB) culture, which was grown for an additional 16 hours at 37 °C with agitation. The starting OD600 of the culture was normalized to 0.1 (or 1.0, as signified^†^ throughout the text, to counteract the bacteriostatic effects of vancomycin and DBCO-vancomycin) in TSB and transferred to an untreated F-bottom 96-well plate. The relevant small molecule or protein was added to the specified final concentrations (typically, 50 or 500 μM for small molecules or 1 μM for the o-mAb protein) in a total volume of 100 μL cell solution and the plate was incubated at 37 °C for 24 hours with agitation. For the OD600 measurements, the bacteria were incubated at 37 °C in an Infinite 200 PRO microplate reader (Tecan Life Sciences, Zürich, Switzerland) and the absorbance at 600 nm was measured every hour.

### Co-incubating *S. aureus* with Cy5-containing secondary labels

After the incubation period, the bacteria were transferred to an untreated V-bottom 96-well plate, spun down at 3,000 RCF for 5 minutes, and washed twice with PBS containing 1 w/v% bovine serum albumin (BSA) (PBS-B). Bacteria were fixed with 2% PFA in PBS for 20 minutes at RT with gentle agitation, then washed twice with PBS-B. Bacteria were resuspended in 50 μL PBS-B optionally containing 1 μM sCy5-azide or 1 μM IgG3-Fc-sCy5-azide (DOL = 1.7) and incubated for 2 hours at RT in the dark with gentle agitation, then washed twice with PBS-B, and analyzed via flow cytometry using a SA3800 Spectral Cell Analyzer (Sony, San Jose, California, USA).

### HEK293T cell culture

HEK293T cells (ATCC, Catalog# CRL-3216) were maintained in DMEM containing 10% FBS and 1% penicillin/streptomycin (a.k.a. complete DMEM) at 37 °C, 5% CO_2_ and passaged upon reaching 70 – 90% confluency. All incubation steps take place at 37 °C, 5% CO_2_ unless otherwise specified.

### Co-incubating HEK293T cells with DBCO-containing small molecules

Cells were seeded at a density of 1.2 x 10^5^ cells/well in a 24-well plate and allowed to re-adhere for 24 hours. The next day, the media was aspirated and 50 or 500 μM of each small molecule in 400 μL complete DMEM was added. The cells were then incubated for 16 hours, washed once with PBS, dissociated using 50 μL trypsin/well for 5 minutes, diluted with 100 μL PBS-B, and transferred to an untreated, U-bottom 96-well plate. The cells were pelleted by centrifuging the plate at 500 RCF for 4 minutes, then resuspended in 50 μL PBS-B containing 1 μM sCy5-azide and incubated on ice in the dark with gentle agitation. After 30 minutes, the cell solutions were diluted with 150 μL PBS-B, then pelleted and resuspended in fresh PBS-B and analyzed via flow cytometry using an SA3800 Spectral Cell Analyzer (Sony, San Jose, California, USA).

### MTS assay

The MTS reagent was prepared by combining 5 mM (2.44 mg/mL) MTS in H_2_O (AAT Bioquest, Catalog# 15710) and phenazine methyl sulfate (PES) powder at a final concentration of 2 mg/mL and 0.21 mg/L, respectively, in Dulbecco’s phosphate buffered saline (DPBS) and adjusting the pH to 6.0 using 1 N HCl. HEK293T cells were seeded at a density of 1.5 x 10^4^ cells/well in a 96-well plate and allowed to re-adhere for 24 hours. The next day, the media was aspirated, 50 or 500 μM of each small molecule in 100 μL complete DMEM was added, and the cells were incubated for 16 hours. Then, 20 uL of the MTS/PES reagent was added to each well and the cells were incubated for an additional 1 hour. The absorbance at 490 nm of each well was measured using an Infinite 200 PRO microplate reader (Tecan Life Sciences, Zürich, Switzerland).

### Labeling IgG3-Fc and Omodenbamab with various small molecules

Human recombinant IgG3-Fc (Sino Biological, Catalog# 13906-HNAH) and human Omodenbamab (MedChemExpress, Catalog# HY-P99770) were desalted into 50 mM borate-buffered saline, pH = 8.4 using a 7 kDa Zeba^TM^ spin desalting column. The proteins were then combined with the specified eq. of sDBCO-PEG_4_-TFP (Vector Laboratories, Catalog# CCT-1527), azide-PEG_4_-NHS (Vector Laboratories, Catalog# CCT-AZ103), sCy5-NHS (Lumiprobe, Catalog# 23320), or sCy5-azide-NHS (Vector Laboratories, Catalog# CCT-1572) and reacted for 1 – 2 hours at RT with end-over-end rotation. Afterwards, each solution was passed through a 7 kDa Zeba^TM^ spin desalting column preloaded with PBS containing 0.05 w/v% Tween 20 (PBS-T) to remove any unreacted material. To obtain an estimate of the protein concentration and DOL, the absorbance at 280 nm and 646 nm (A_280_ and A_646_) were measured using a NanoDrop 2000c spectrophotometer with a 10 mm, 50 μL UV/VIS cuvette (Eppendorf, Catalog# 952010051). For IgG3-Fc labeled with sCy5-azide-NHS, the concentration in μM was calculated using the equation (A_280_-(A_646_*0.04))/40,560*10^6^. For Omodenbamab labeled with sCy5-NHS, the concentration in μM was calculated using the equation (A_280_-(A_646_*0.04))/210,000*10^6^. The DOL for both proteins was estimated using the equation A_646_/(271,000*conc. [μM])*10^6^.

### SDS-PAGE gel

For the storage stability studies, IgG3-Fc labeled with either sDBCO-PEG_4_-TFP or azide-PEG_4_-NHS, or unlabeled (DMSO vehicle control) were stored at 4 °C, RT, or 37 °C for 72 hours prior to running on a gel.

For the high-fidelity estimates of the DOL following labeling with sCy5-azide-NHS, 4 μg of the IgG3-Fc (unlabeled) or IgG3-Fc-sCy5-azide protein in PBS-T were combined with 20 eq. of 5 kDa DBCO-mPEG (Vector Laboratories, Catalog# CCT-A118) and reacted overnight at RT.

As appropriate, samples were combined with 4X Laemmli sample buffer (reducing), heated to 95 °C for 5 minutes, and loaded onto a 4-20% Mini-PROTEAN TGX precast SDS-PAGE gel (Bio-Rad, Catalog# 4561094) alongside a protein ladder and ran at 200 V for 30 minutes. The gel was rinsed 3 times with deionized H_2_O and stained overnight using rapid Coomassie stain (RPI, Catalog# RCS-50) with 1 mL of 20 w/v% NaCl added per 10 mL of staining solution. The following day, the gel was rinsed 3 times with deionized H_2_O prior to imaging. Band intensity was quantified using ImageJ.

### CD64 ELISA

NUNC MaxiSorp ELISA plates were coated with 0.5 μg/mL human CD64 (Sino Biological, Catalog# 10256-H08H) in 50 mM carbonate-bicarbonate buffer, pH = 9.6, and incubated overnight at 4 °C with agitation. The next day, the plates were washed 3 times with PBS-T and blocked with 300 μL 5% nonfat milk in PBS-T for 1 – 2 hours at RT with agitation. The wells were emptied by inverting and shaking the plates and samples diluted in 0.5% nonfat milk in PBS-T (diluent) were added and the plates were incubated for 1 – 2 hours at RT with agitation. The plates were then washed 3 times with PBS-T and 100 μL of 0.25 μg/mL goat anti-human IgG (H+L)-HRP secondary (Novus Biologicals, Catalog# NBP1-74957) in diluent was added. The plate was additionally incubated for 1 – 2 hours at RT with agitation. Finally, the plate was washed 5 times with PBS-T and 100 μL KPL TMB peroxidase substrate (SeraCare, Catalog# 5120-0047) was added and the plate was allowed to develop for 5 minutes at RT with agitation. The reaction was halted by adding 50 μL 0.5 M and the absorbance was read at 450 and 600 nm immediately afterwards using an Infinite 200 PRO microplate reader (Tecan Life Sciences, Zürich, Switzerland).

## ACKNOWLEDGEMENTS

We’d like to thank the lab of Dr. Katherine Lemon at BCM for donating the *S. aureus* Newman that was used in this work. We would also like to acknowledge the support of the Shared Equipment Authority at Rice University.

## Notes

### Competing Interest Statement

The authors have declared no competing interest.

