## Supplemental Figures for "Novel approaches to label the surface of *S. aureus* with DBCO for click chemistry-mediated deposition of sensitive cargo"

### SUPPORTING INFORMATION

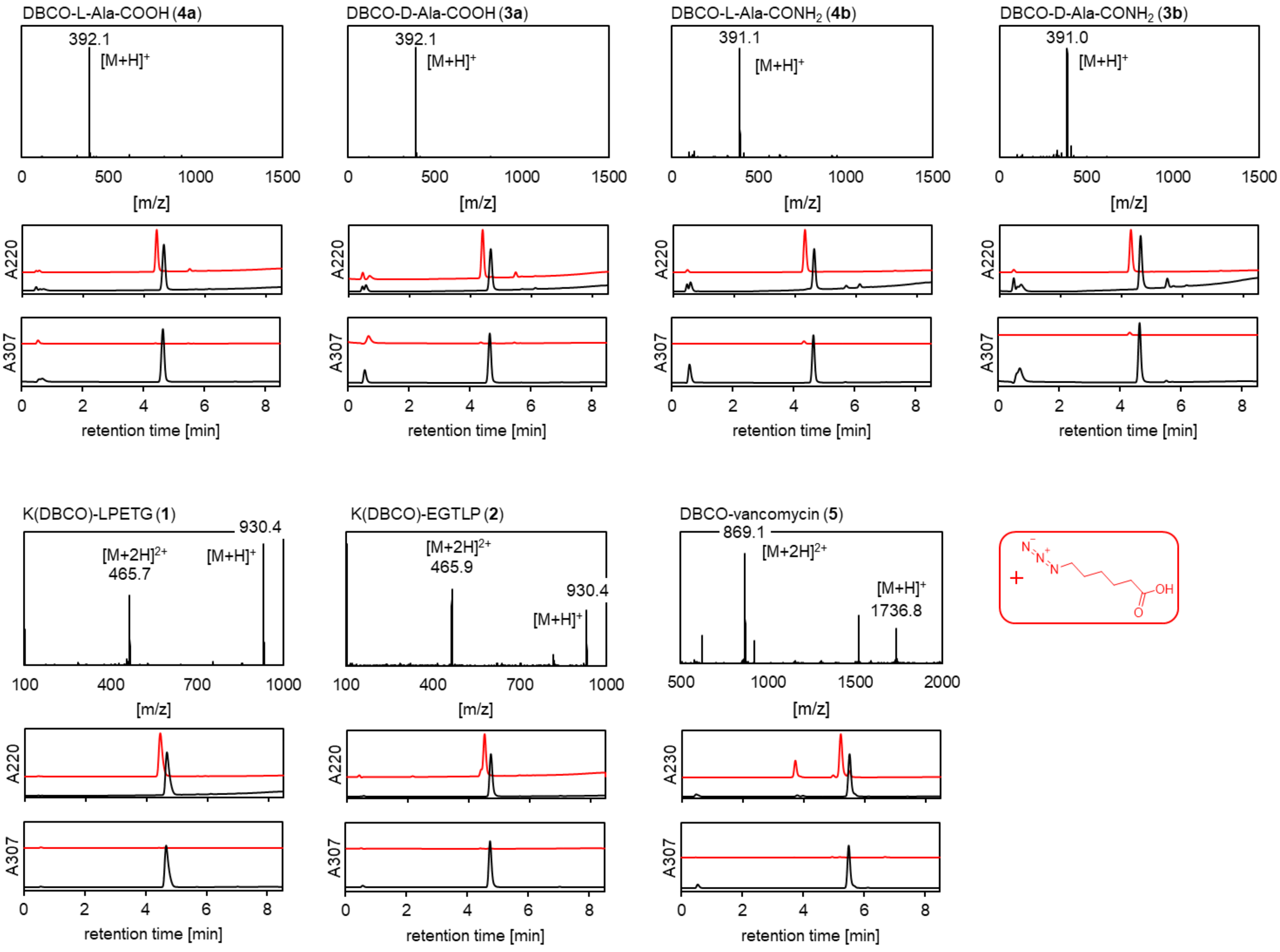


**Figure S1 | Characterization of the small molecules used in this work.** All small molecules were analyzed via MS-ESI and RP-UPLC. Each was additionally reacted with a large excess of 6-azido hexanoic acid and reanalyzed via RP-UPLC to demonstrate a shift in the peak at 220/230 nm and ablation of the peak at 307 nm, signifying complete formation of the azide-DBCO complex.

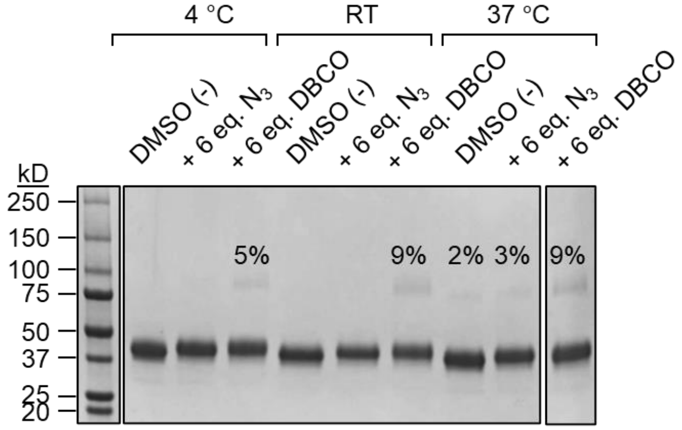


**Figure S2 | Stability of IgG3-Fc protein labeled with azido vs. DBCO groups.** 4 μg IgG3-Fc labeled with either 6 molar equivalents (eq.) of sDBCO-PEG$_4$-TFP or azide-PEG$_4$-NHS, or unlabeled (DMSO vehicle control) were stored at 4 °C, RT, or 37 °C for 72 hours and ran on an SDS-PAGE gel. The relative band intensity was quantified via ImageJ. The unlabeled protein sample and the sample labeled with azide produced two bands (one having a higher MW) when both samples were stored at 37 °C, comprising 2 – 3% of the total protein content in either case but appeared stable when stored at 4 °C and RT. In comparison, the protein sample labeled with DBCO produced a second band at all three storage temperatures, comprising 5 – 9% of the total protein content and signifying an increase in aggregation propensity.

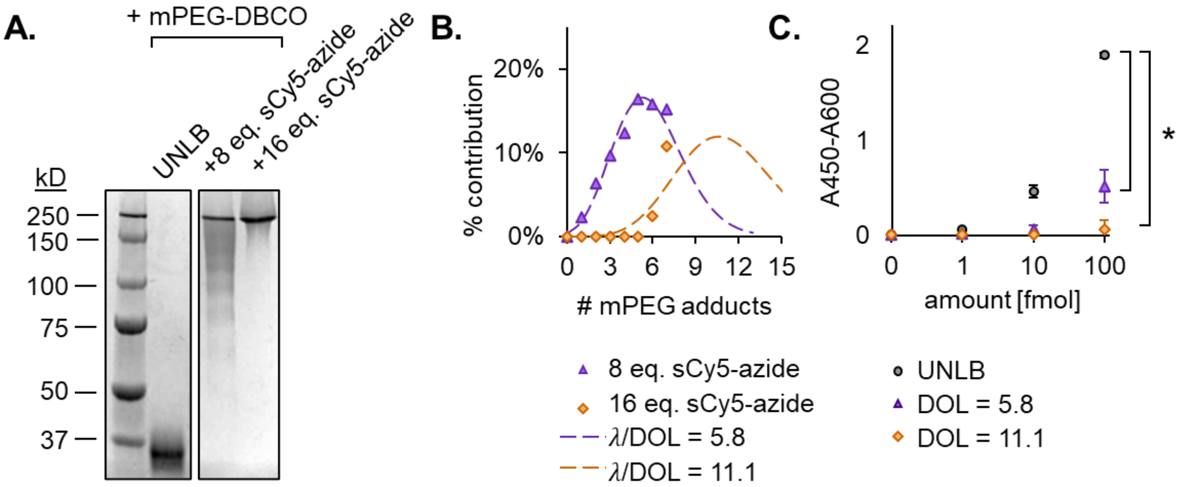


**Figure S3 | Labeling of IgG3-Fc with high equivalencies of azido groups and assessing protein conjugate characteristics.** Human IgG3 was labeled with sCy5-azide-NHS as in Figure 5 and (**A**) run on an SDS-PAGE gel to determine the degree of labeling (DOL). (**B**) The intensity of each adduct band in (**A**) was quantified via ImageJ and a Poisson curve was fit to the resulting data distribution to yield the λ parameter, which is synonymous with the DOL. (**C**) A sandwich ELISA was performed to determine whether the IgG3-Fc-sCy5-azide retained the ability to bind CD64 with *$p < 0.05$. Along with the data in Figure 5, these results demonstrate that an increase in sCy5-azide DOL tends to result in a commensurate decrease in the ability of IgG3-Fc to bind CD64.
